# MicroRNAs alteration and unique distribution in the soma and synapses of substantia nigra in Parkinson’s disease

**DOI:** 10.1101/2025.03.12.642888

**Authors:** Morgan N. Ogwo, Gunjan Goyal, Peter Zotor, Bhupender Sharma, Daniela Rodarte, Rajkumar Lakshmanaswamy, Subodh Kumar

**Author notes:** Correspondence: **Subodh Kumar, PhD,** Assistant Professor, Center of Emphasis in Neuroscience, Department of Molecular and Translational Medicine, Texas Tech University Health Sciences Center El Paso, 130 Rick Francis, El Paso, Texas, USA 79905.

## Abstract

Parkinson’s disease (PD) is the second most common neurodegenerative condition after Alzheimer’s. Abnormal accumulation of alpha-synuclein (α-syn) aggregates disrupts the balance of dopaminergic (DA-ergic) synapse components, interfering with dopamine transmission and leading to synaptic dysfunction and neuronal loss in PD. However exact molecular mechanism underlying DA-ergic neuronal cell loss in the SNpc in not known. MicroRNAs (miRNAs) are observed in various compartments of neural elements including cell bodies, nerve terminals, mitochondria, synaptic vesicles and synaptosomes. However, miRNAs expression and cellular distribution are unknown in the soma and synapse compartment in PD and healthy state. To address this void of information, we isolated synaptosomes and cytosolic fractions (soma) from post-mortem brains of PD-affected individuals and unaffected controls (UC) and processed for miRNA sequencing analysis. A group of miRNAs were significantly altered (*p* < 0.05) with high fold changes (variance +/− > 2-fold) in their expressions in different comparisons: 1. UC synaptosome vs UC cytosol, 2. PD synaptosome vs PD cytosol, 3. PD synaptosome vs UC synaptosome, 4. PD cytosol vs UC cytosol. Our study unveiled some potential miRNAs in PD and their alteration and unique distribution in the soma and synapses of SNpc in PD and controls. Further, gene ontology enrichment analysis showed the involvement of deregulated miRNAs in several molecular function and cellular components: synapse assembly formation, cell junction organization, cell projections, mitochondria, Calcium ion binding and protein binding activities.

## INTRODUCTION

Parkinson’s disease (PD) is a neurodegenerative disorder characterized by progressive motor impairment, autonomic dysfunction, and neuropsychiatric symptoms. Since first being described in the literature as “paralysis agitans” by Dr. James Parkinson in 1817, PD has continued to befuddle the medical and academic community. According to the Global Burden of Disease (GBD) study, there were 6.1 million individuals living with PD in 2016, a 21.7% increase in the prevalence since 1990 (Ou et al., 2021). Although the average lifespan of new cases until rehabilitation or death has increased (Ou et al., 2021), to date, a cure has not been documented and an exact cause remains elusive. However, research has elucidated a few strong associations and hypotheses have been established regarding the etiology and pathogenesis of PD.

Synapse dysfunction, or poor pre-synaptic and post-synaptic functionality, has emerged as a player in the pathogenesis and progression of PD. This concept began with post-mortem microscopic and histopathological analyses of PD-affected individuals, which led to formation of hypotheses such as *Braak’s Hypothesis*, the *Body First vs Brain First Hypothesis*, and the *Dying Back Hypothesis* which surmise that some event occurs distally in the nervous system that triggers some form of retrograde axonal transport or neurodegeneration eventually reaching the regions of the brain (e.g. substantia nigra) associated with classical symptoms of PD. Basically, the neurodegeneration observed in PD begins with a toxic insult at the synapse. These hypotheses are supported by a multitude of research findings detailing dopaminergic synaptic terminal defects and axonal degeneration being observed prior to cell body loss (Herkenham et al., 1999; Serra et al., 2002; Meissner et al., 2003; Li et al., 2009; Chu et al., 2012; Morales et al., 2021).

Several questions can be posed from these findings. If the neurodegeneration in PD starts with some sort of damage at the synapse, what exactly is the insult and how can its effects be modulated? PD is commonly recognized as a condition that occurs following the significant loss of midbrain dopaminergic neurons. Current therapeutic standards primarily involve replacement of dopamine using levodopa, stimulating remaining dopamine receptors using dopamine receptor agonists, and reducing dopamine metabolism using a group of enzyme inhibitors. However, there are no agents that have been shown to prevent, reduce the progression of, or cure PD. Evidence has pointed to genetic implications, mitochondrial factors, and impairment in synaptic plasticity among other factors. One of the factors that is less studied is the role microRNA (miRNA) play in synapse dysfunction especially as it relates to PD.

Synapses are a key component of healthy brain function. Functionally, they are the connection point for neurons and consist of the pre-synaptic compartment, post-synaptic compartment, and the space between, in which neurotransmitters and other substances are transferred. Synapse integrity (functions, number, and structure) is vital for effective and efficient neurotransmission as well as to meet the complex demands of the nervous system. Synapse components can actually be extracted from post-mortem brain tissue in an intact form referred to as “synaptosomes or synapto-neurosomes”. Synaptosomes are one of the greatest means by which synapse dysfunction can be studied in neurodegenerative diseases. It was synaptosome technology that was utilized by Kahle and Colleagues to demonstrate that alpha-synuclein ([]-Syn) localizes to the synapse back in 2000 (Kahle et al., 2000). In PD, the synaptosome functionality is impaired by aberrant accumulation of alpha-synuclein ([]-Syn) (Desplats et al., 2009; Volpicelli et al., 2011; Lindestam Arlehamn et al., 2020). The disruption of the integrity of synapses in PD is observed in impaired neurotransmitter release, compromised synaptic mitochondrial function, disorganized synaptic vesicle trafficking, release, and recycling, inadequate receptor and transporter distribution, stifled synaptic plasticity, and inflammation among other things (Centonze et al., 1999; Calabresi et al., 2000; Calabresi et al., 2007; Li et al., 2010; Volpicelli et al., 2011; Rockenstein et al., 2014; Matikainen-Ankney et al., 2016; Lindestam Arlehamn et al., 2020; Henrich et al., 2023; Masato et al., 2023). What’s more interesting is that miRNA may regulate all of these functions.

MiRNAs are small non-coding RNA molecules with gene regulatory properties. Some miRNAs are localized to subcellular compartments including endoplasmic reticula, endosomes, golgi apparatuses, lysosomes, mitochondria, and multivesicular bodies (O’brien et al., 2018). Interestingly, they are also localized to the synapse and can be found in synaptosomal fractions playing a role in local protein synthesis (Lugli et al., 2008; Xu et al., 2013; Li et al., 2015; Boese et al., 2016). These miRNAs, often referred to in the literature as “enriched at the synapse” herein referred to as synaptic miRNA or synaptosomal miRNA, may occupy a key role in regulating the integrity of the synapse. Several studies showed these miRNAs are involved in affecting synaptic structure (Edbauer et al., 2010; Lee et al., 2016; Tsujimura et al., 2016; Wu et al, 2018; Mcneill et al., 2020). Interestingly, there are those that have been identified as brain-specific and are involved in a host of functions including neurogenesis, synaptic plasticity, and neuroinflammation (Sempere et al., 2004; Makeyev et al., 2007; Ponomarev et al., 2011).

There is no doubt that synaptic miRNAs are involved in a variety of synaptic functions. However, the role of synaptosome-specific miRNAs is not yet determined in the progression of PD. Very few, if any, published reports exist regarding the roles synaptosome-specific miRNAs play in PD. Furthermore, it is unclear whether the variety, distribution, and functions of synaptosomal miRNAs differs from cytosolic miRNAs. This study sought to unravel this information. In turn, synaptosomal and cytosolic miRNA were identified, classified and compared. Additionally, potential molecular and biochemical associations between these miRNAs and the progression of PD were established. Our study addressed four important research questions: (1) Are miRNA(s) levels altered at the synaptosome in PD? (2) If so, are synapse miRNAs expressed differently in PD than in a healthy state? (3) Are synaptosomal miRNAs expressed differentially in the cytosol? and (4) What function do synaptosomal miRNAs play in synaptic activity and neurotransmission in PD? Current study unveiled some specific miRNAs in PD and their alteration and unique distribution in the soma and synapses of SNpc in PD and controls. Overall, the focus of this study is to discover synaptosomal miRNAs and understand their positive and negative roles in PD progression.

## MATERIALS AND METHODS

### PD and unaffected controls post-mortem brain samples

Postmortem brain samples from PD patients and unaffected controls were obtained from NIH NeuroBioBanks-(1) Harvard Brain Tissue Resource Center (HBTRC), Mclean Hospital, Mail Stop 138, 115 Mill Street, Belmont, MA 02478-1064. (2) Brain Endowment Bank, University of Miami, Millar School of Medicine, 1951, NW 7th Avenue Suite 240, Miami, FL. (3) University of Maryland Brain and Tissue Bank (UMBTB), University of Maryland School of Medicine, Department of Pediatrics, 655 W. Baltimore St., 13-013 BRB, Baltimore, MD 21201-1559. Brain tissues were dissected from the substantia nigra of PD patients (n = 5) and age and sex-matched unaffected controls (UC) (n = 5). Demographic and clinical details of study specimens are provided in Table 1. The study was conducted at the Molecular and Translational Medicine Department, Texas Tech University Health Sciences Center El Paso, and the Institutional Biosafety Committee (IBC protocol # 22008) approved the study protocol for the use of human postmortem brain tissues obtained from NIH NeuroBioBanks. The NIH NeuroBioBanks mentioned above operated under their institution’s IRB approval, and they obtained written informed consent from the donors (Kumar et al., 2018; Kumar et al., 2022).

**Table 1.** Details of PD and unaffected control post-mortem brains.

| 1. Harvard Brain Tissue Resource Center (HBTRC) |  |  |  |  |  |  |  |
| --- | --- | --- | --- | --- | --- | --- | --- |
| Sample | HSB | Age | Sex | Race | Clinical Brain Diagnosis | Brain region | PMI (hours) |
| PD 1 | S15419 | 73 | F | Unknown | PD | Substantia nigra | 20.97 |
| PD 2 | S01706 | 37 | M | White | PD | Substantia nigra | 22.62 |
| PD 3 | S05845 | 76 | F | White | PD | Substantia nigra | 22.63 |
| PD 4 | S12782 | 72 | M | Unknown | PD | Substantia nigra | 19.2 |
| PD 5 | S19897 | 70 | F | White | PD | Substantia nigra | 19.58 |
| 2. Brain Endowment Bank, University of Miami |  |  |  |  |  |  |  |
| Sample | Tissue Code | Age | Sex | Race | Clinical Brain Diagnosis | Brain region | PMI (hours) |

| UC 1 | HCTYH A017 | 71 | F | White | None | Substantia nigra | 16.4 |
| --- | --- | --- | --- | --- | --- | --- | --- |
| UC 2 | HCTYD A034 | 68 | F | White | None | Substantia nigra | 19.05 |
| UC 3 | HCTZZ JA008 | 70 | M | White | None | Substantia nigra | 27.2 |
| 3. University of Maryland Brain and Tissue Bank (UMBTB) |  |  |  |  |  |  |  |
| Sample | UMBN# | Age | Sex | Race | Clinical Brain Diagnosis | Brain region | PMI (hours) |
| UC 4 | 1376 | 37 | M | Black | None | Substantia nigra | 12 |
| UC 5 | 5618 | 68 | F | White | None | Substantia nigra | 24 |

### Synaptosome extraction

Synaptosomes were extracted using Syn-PER Reagent as described in our previous study (Kumar et al., 2022). Briefly, 50 mg of brain tissue was used from each sample for synaptosome extraction in 1 ml of Syn-PER Reagent. Tissues were homogenized slowly by Dounce glass homogenization on ice with ∼10 slow strokes. Samples were centrifuged at 1400 g for 10 minutes at 4 °C to remove the leftover tissue debris. After centrifugation, the supernatant was transferred to a new tube. Again, the supernatant (homogenate) was centrifuged at a speed of 15,000 g for 20 min at 4 °C. The supernatant was removed as a cytosolic fraction, and synaptosomes were recovered in pellet form. The synaptosome pellets were processed for RNA extraction.

### RNA extraction from soma and synaptosome

Total RNA, including miRNAs, was extracted from the synaptosomes of PD and control postmortem brain samples using the TriZol reagent with some modifications. RNA quality and purity were determined by NanoDrop and equal amount of RNA from all twenty sample: PD cytosolic fraction (soma) (n=5), PD synaptosome fraction (n=5), UC cytosolic fraction (soma) (n=5) and UC synaptosome fraction (n=5) were processed for miRNA sequencing analysis following previous lab study (Kumar et al., 2022; Kumar et al., 2025).

### MiRNA sequencing analysis

The miRNAs HiSeq analysis were performed commercially at LC Sciences Houston, Texas following our previous lab publications (Kumar et al., 2022; Kumar et al., 2025). MiRNA HiSeq analysis flow chart is shown in the Supplementary Information Figure 1 (SI Figure 1). All extracted RNA was used in the library preparation following Illumina’s TruSeq -small-RNA-sample preparation protocols (Illumina, San Diego, CA, USA). Quality control analysis and quantification of the DNA library were performed using Agilent Technologies 2100 Bioanalyzer High Sensitivity DNA Chip. Single-end sequencing 50bp was performed on Illumina’s Hiseq 2500 sequencing system following the manufacturer’s recommended protocols.

**Figure 1.**
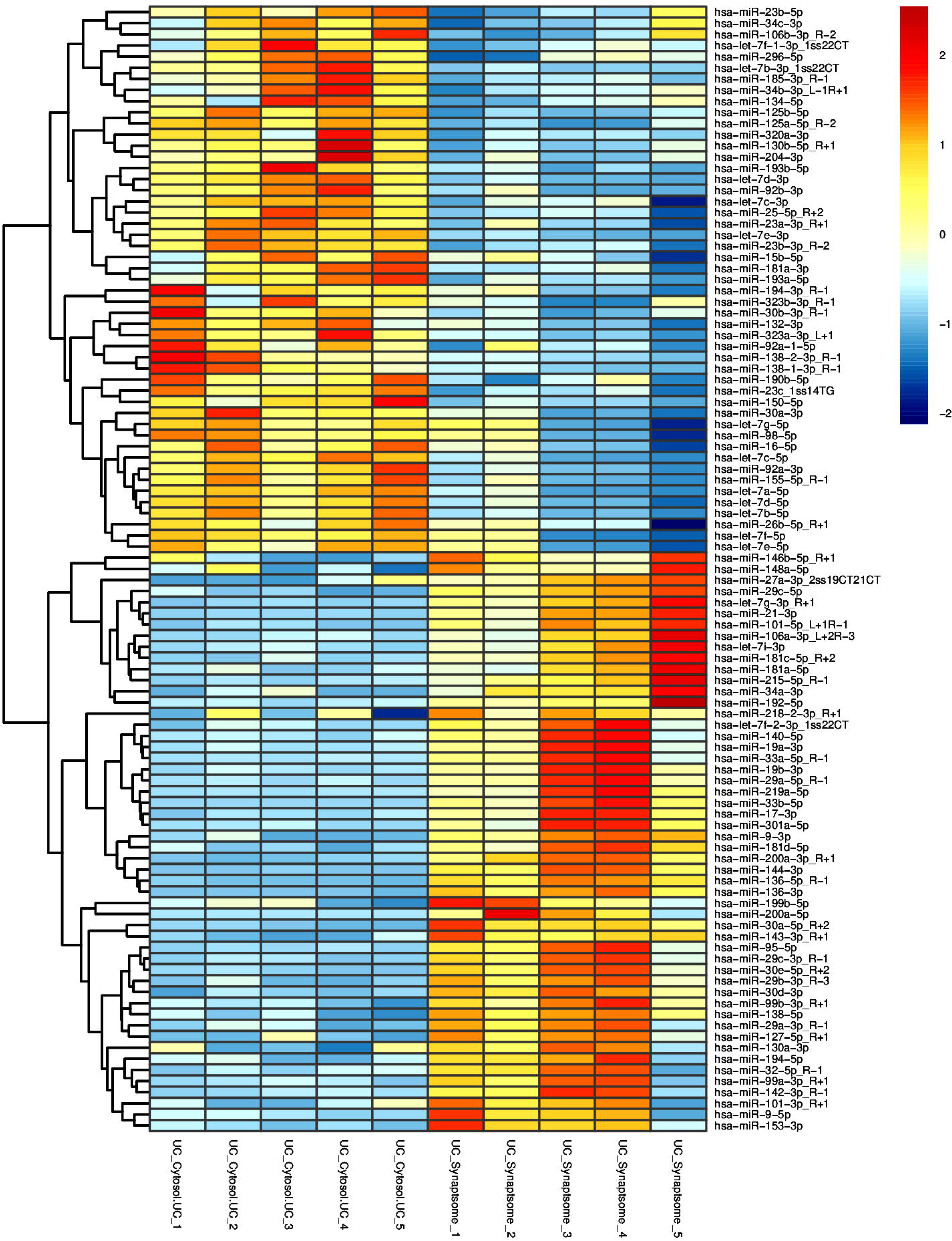
MiRNAs expression in UC synaptosome vs UC cytosol. Hierarchical clustering and heatmap of 101 significantly distributed miRNAs in synaptosome and cytosol in UC samples. Red color intensity showed the miRNAs upregulation and blue color intensity showed the miRNAs downregulation.

Raw reads were subjected to an in-house program, ACGT101-miR (LC Sciences, Houston, Texas, USA) to remove adapter dimers, junk, low complexity, common RNA families (rRNA, tRNA, snRNA, snoRNA) and repeats. Subsequently, unique sequences with length in 18∼26 nucleotide were mapped to specific species precursors in miRBase 22.0 by BLAST search to identify known miRNAs and novel 3p- and 5p-derived miRNAs. Length variation at 3’ and 5’ ends and one mismatch inside the sequence was allowed in the alignment. The unique sequences mapping to specific species of mature miRNAs in hairpin arms were identified as known miRNAs. The unique sequences mapping to the other arm of known specific species precursor hairpin opposite to the annotated mature miRNA-containing arm were considered novel 5p- or 3pderived miRNA candidates. The remaining sequences were mapped to other selected species precursors (with the exclusion of specific species) in miRBase 22.0 by BLAST search, and the mapped pre-miRNAs were further BLASTed against the specific species genomes to determine their genomic locations. The unmapped sequences were BLASTed against the specific genomes, and the hairpin RNA structures containing sequences were predicated from the flank 80 nt sequences using RNAfold software (http://rna.tbi.univie.ac.at/cgi-bin/RNAfold.cgi). The criteria for secondary structure prediction were: (1) number of nucleotides in one bulge in the stem (≤12) (2) number of base pairs in the stem region of the predicted hairpin (≥16) (3) cutoff of free energy (kCal/mol ≤-15) (4) length of hairpin (up and down stems + terminal loop ≥50) (5) length of hairpin loop (≤20). (6) number of nucleotides in one bulge in mature region (≤8) (7) number of biased errors in one bulge in mature region (≤4) (8) number of biased bulges in mature region (≤2) (9) number of errors in mature region (≤7) (10) number of base pairs in the mature region of the predicted hairpin (≥12) (11) percent of mature in stem (≥80). Differential expression of miRNAs based on normalized deep-sequencing counts was analyzed selectively using the Fisher exact test, Chi-squared 2X2 test, Chi-squared nXn test, Student t test, or ANOVA, depending on the experiment’s design. The significance threshold was set to 0.01 and 0.05 in each test.

### Bioinformatics analysis

To predict the genes targeted by the most abundant miRNAs, two computational target prediction algorithms (TargetScan 50 and Miranda 3.3a) were used to identify miRNA binding sites. Finally, the data predicted by both algorithms were combined, and the overlaps were calculated. The GO terms and KEGG Pathway of these most abundant miRNAs and miRNA targets were also annotated.

Biotype classification and composition analyses were done using the Ensembl database (REST API) on filtered differentially expressed genes. Gene Ontology (GO) was carried out by querying the bioinformatic Database for Annotation, Visualization, and Integrated Discovery (DAVID). The following annotation categories were included in the analysis: KEGG Pathways, GO: Biological Processes, GO: Molecular Function, GO: Cellular Components. The data was visualized in R statistical software (version 4.0.3) using the packages ggplot2 and plotly/heatmap for dot plots and heatmaps, respectively.

### Statistical analysis

The statistical analyses of biological data were conducted using the student’s *t*-test to analyze two groups of samples: control *versus* PD miRNAs for both synaptosome and cytosol analysis. Statistical parameters were calculated using GraphPad Prism software, v6 (GraphPad, San Diego, CA, USA) (www.graphpad.com). The *P* values < 0.05 were considered statistically significant.

## RESULTS

### MicroRNA expression in unaffected control synaptosome versus soma

The miRNA HiSeq data of synaptosomal and cytosolic fractions (soma) showed a total of 294 mature miRNAs were found to be significantly differentially distributed between UC synaptosomal and UC cytosolic fractions (Supplementary Table 1). Of the 294, 30 were significantly altered at a p-value < 0.001, 80 at a p-value < 0.01, and 184 at a p-value < 0.05. Only 104 (35.4%) were highly expressed, 130 (44.2%) were upregulated whilst 164 (55.8%) were downregulated. Heat map showed most significantly deregulated 101 miRNA in UC cytosol vs UC synaptosome, expression of 49 miRNAs were more in cytosol while 52 miRNAs were highly expressed in synaptosome as shown in Figure 1. Notably, miRNAs hsa-miR-143-3p_R+1 and hsa-miR-138-5p were significantly upregulated while hsa-miR-125a-5p_R-2 and hsa-miR-23b-3p_R-2 were downregulated as shown in the volcano plot in Figure 2A. Figure 2B, showed the miRNAs correlation and distribution in UC cytosol and UC synaptosome, based on their log10 mean value. Figure 2C showed a significant overlap of expression of miRNAs with some differential localization of miRNAs between synaptosome and cytosol. Further, KEGG/GO enrichment analysis showed that involvement of miRNAs in top 20 cellular components and biological pathways (Figure 2D). These observations reveal a high variability between the expression of specific miRNA at the synapse versus in other compartments of the neural tissue and suggest some sort of importance and perhaps difference in demand/function with those that are highly expressed. Interestingly, 68 (65.4%) of the miRNA that were highly expressed were downregulated in the synapse. Perhaps in non-PD affected individuals, these miRNAs have higher demand, given their various functions, within other compartments of neurons than in the synapses. MiRNAs were characterized on several selection criteria *p*-values, fold change, standard deviation, expression priority, transcript ID, chromosome location, strand specificity, start and stop codon, and validated gene symbols. Gene filter analysis of the total miRNA pool shows only 12.7% of the miRNA population showed a significant difference in the synaptosomal compartment in comparison to the cytosolic compartment. Only 5.62% of the population was significantly upregulated and 7.18% were significantly downregulated.

**Figure 2.**
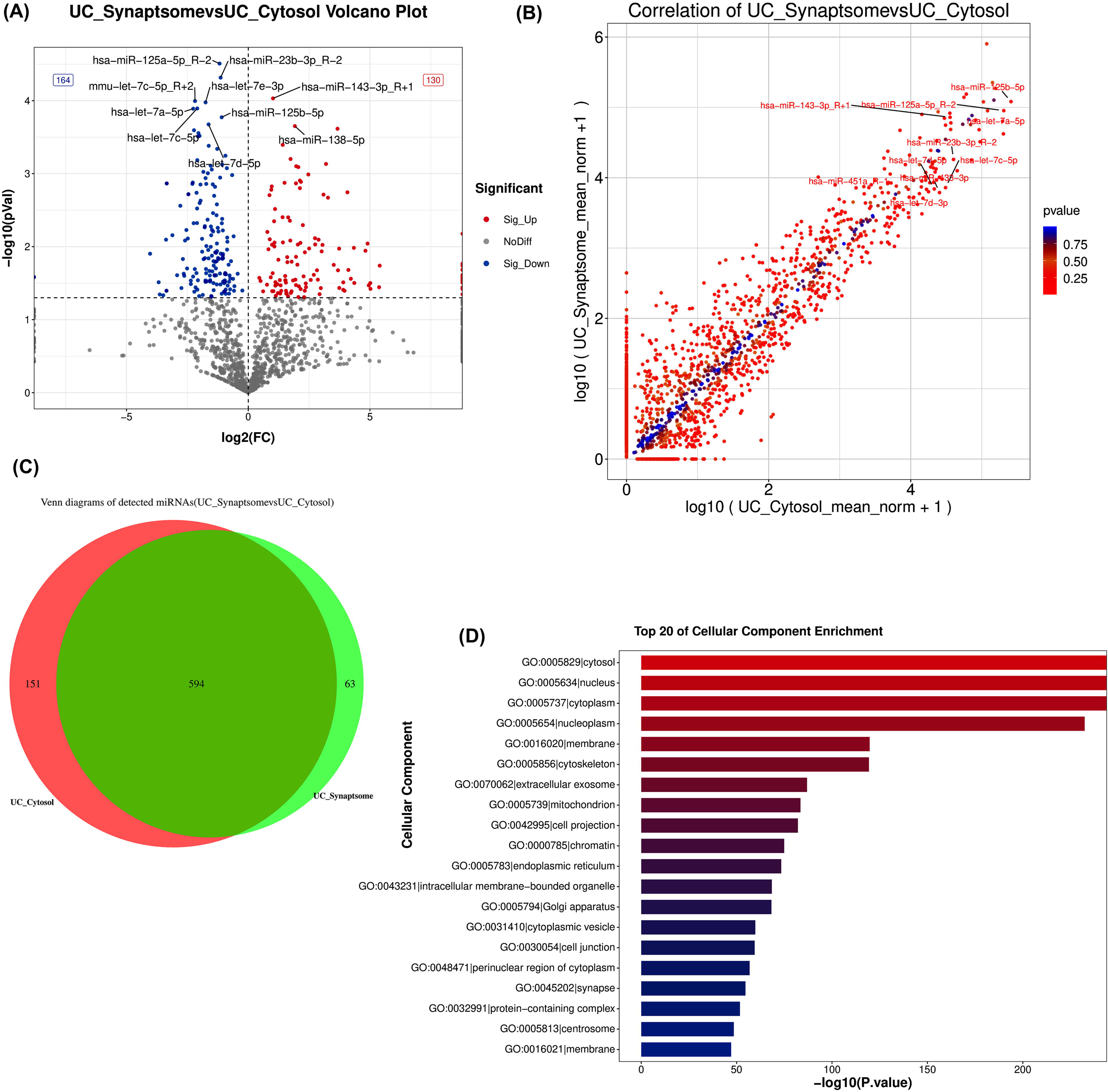
MiRNAs expression in UC synaptosome vs UC cytosol. **(A)** Volcano plot depicting differentially expressed miRNAs in UC synaptosome vs UC cytosol. Red dot showed the miRNAs upregulation, blue dot showed the miRNAs downregulation, and gray dot showed no difference. Significantly upregulated and downregulated miRNAs are highlighted, illustrating distinct expression profiles between the two cellular compartments. **(B)** Correlation plot comparing miRNA expression levels between UC synaptosome and UC cytosol. Each dot corresponds to an individual miRNA, with the diagonal line indicating equal expression between the two compartments. Deviations from the diagonal reflect differential distribution of miRNAs between the synaptosome and cytosol with red color intensity indicating level of significance. **(C)** Venn diagram illustrating the distribution of miRNAs between UC synaptosome and UC cytosol with significant overlap between the two compartments shaded in deep green color, and compartment specific expression of miRNAs, shaded in light green color for synaptosome and shaded in red color for cytosol. **(D)** Bar chart depicting the top 20 cellular component localization of miRNAs showing differential enrichment of miRNAs across cellular regions.

### MicroRNAs expression in PD synaptosomes versus PD soma

Next, we analyzed the microarray data for changes in miRNA expression in PD synaptosomal fractions vs PD cytosol fractions. A total of 202 mature miRNA were determined to be significantly differentially distributed between the PD synaptosomal and PD cytosolic fractions (Supplementary Table 2). Of the 202, 10 were significantly altered at a p-value < 0.001, 61 at a p-value < 0.01, and 131 at a p-value < 0.05. Only 40 (19.8.%) were highly expressed 86 (42.6%) were upregulated whilst 116 (57.4%) were downregulated. Heat map showed most significantly deregulated 101 miRNA in PD cytosol vs PD synaptosome, expression of 54 miRNAs were more in cytosol while 47 miRNAs were highly expressed in synaptosome as shown in Figure 3. In particular, miRNAs hsa-miR-30b-3p_R-1 and hsa-miR-33a-3p_R-1 were significantly upregulated while hsa-miR-6507-5p, hsa-let-7e-3p, and hsa-miR-30b-3p_R-1 were downregulated as shown in the volcano plot in Figure 4A. The correlation plot in Figure 4B showed similar expression of miRNAs in the synaptosome and cytosol. Figure 4C showed a significant overlap of expression of miRNAs with some differential localization of miRNAs between synaptosome and cytosol. In terms of molecular function enrichment, these miRNAs were significantly involved in protein binding and less involved in protein kinase binding (Figure 4D). Like seen in non-PD affected individuals, there appears to be a high variability between the expression of miRNA at the synapse versus in other compartments of the neural tissue. However, there are much less highly expressed miRNA in this cohort. Interestingly, 30 (75%) of the miRNA that were highly expressed were downregulated in the synapse. It is possible that the PD pathology plays some role in the observed high percentage of downregulated miRNA that are usually highly expressed in the synaptic compartment vs the cytosolic. This finding may warrant further exploration. Gene filter analysis of the total miRNA pool shows only 6.3% of the miRNA population showed a significant difference in the synaptosomal compartment in comparison to the cytosolic compartment. Only 2.70% of the population was significantly upregulated and 3.64% were significantly downregulated.

**Figure 3.**
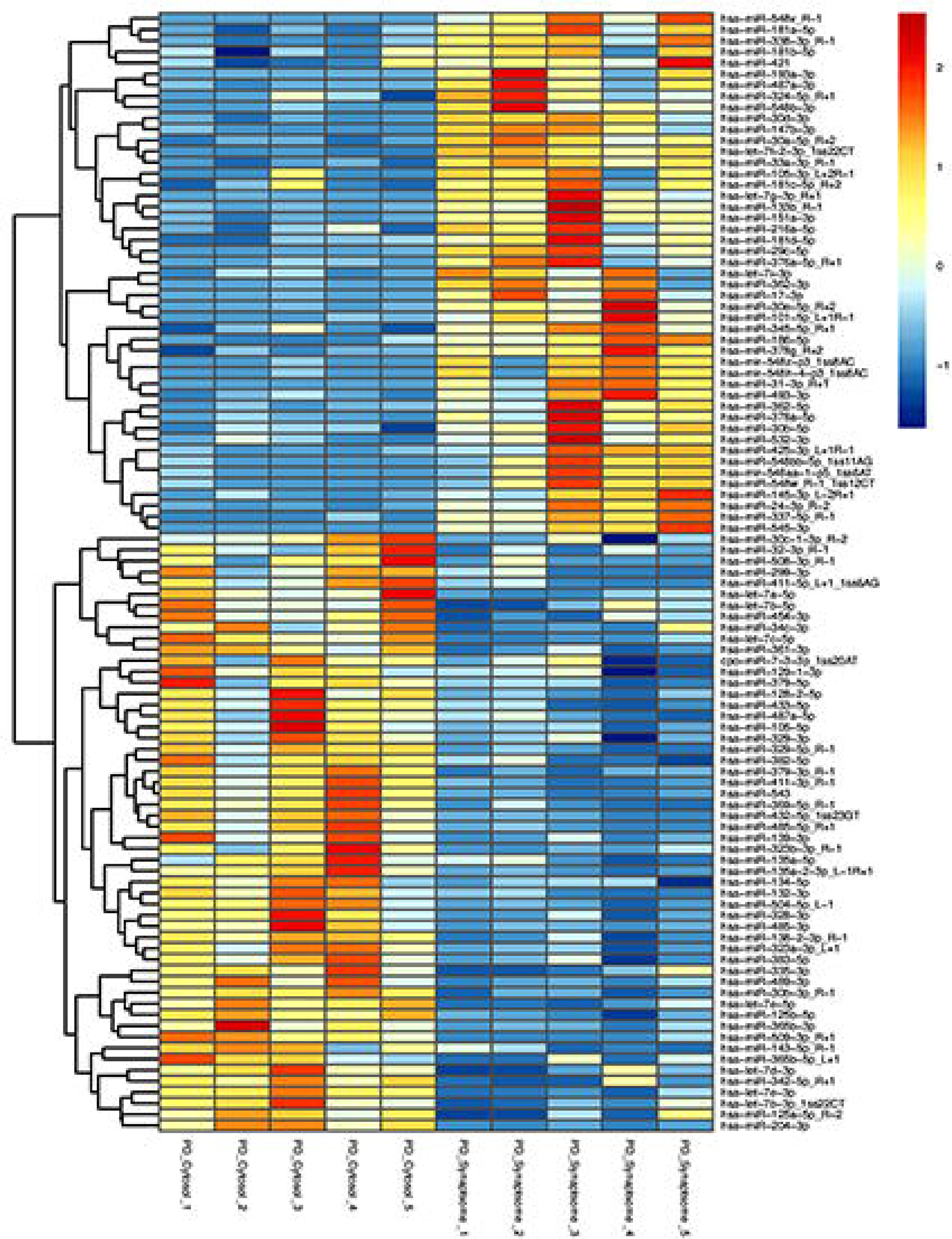
MiRNAs expression in PD synaptosome versus PD cytosol. Hierarchical clustering and heatmap of 101 significantly distributed miRNAs synaptosome and cytosol in PD samples. Red color intensity showed the miRNAs upregulation and blue color intensity showed the miRNAs downregulation.

**Figure 4.**
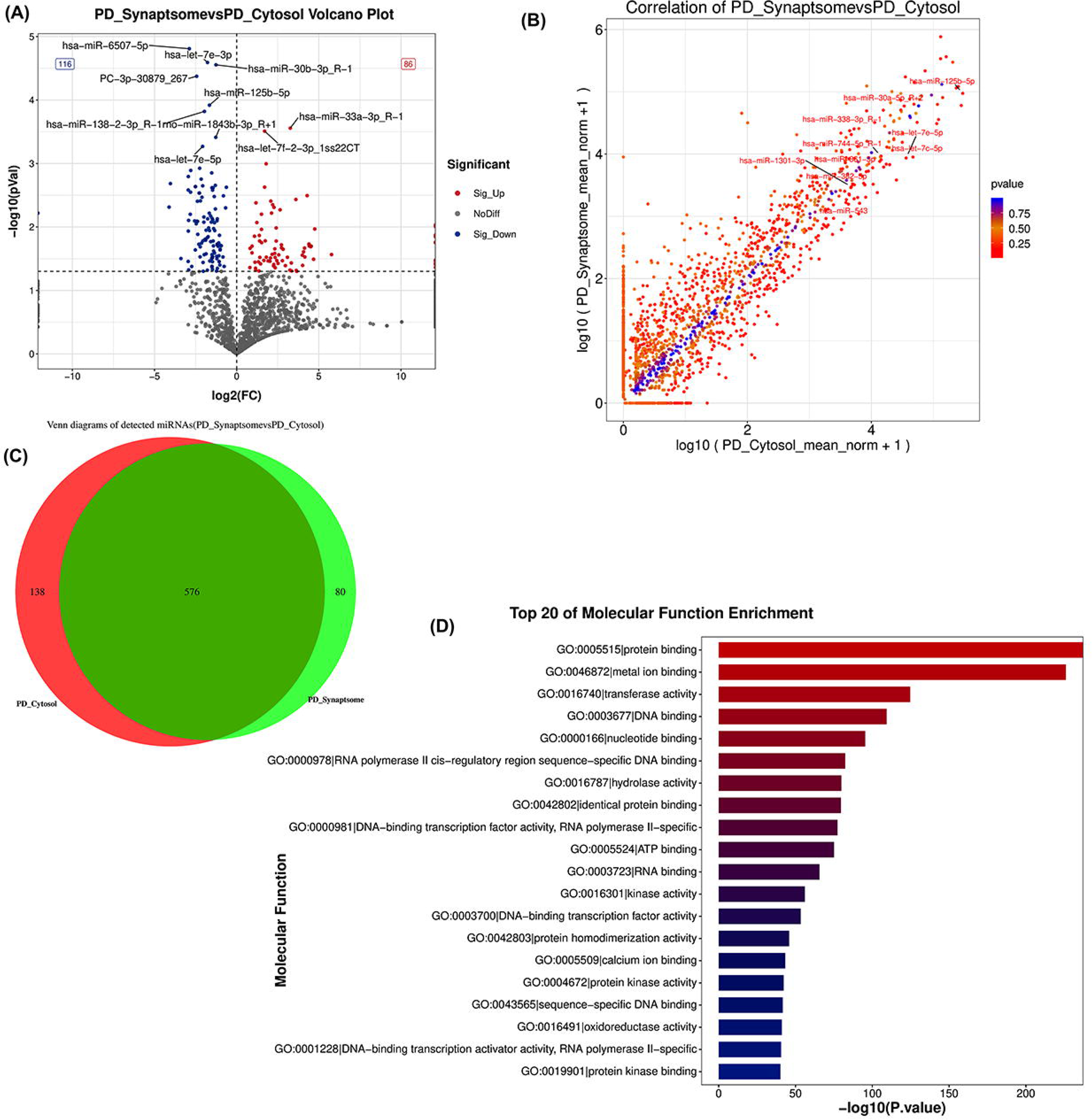
MiRNAs expression in PD synaptosome versus PD cytosol. **(A)** Volcano plot depicting differentially expressed miRNAs in PD synaptosome vs PD cytosol. Red dots show miRNAs upregulation, blue dots show miRNAs downregulation, and gray dots show no significant difference. Significantly upregulated and downregulated microRNAs are highlighted, illustrating distinct expression profiles between the two cellular compartments. **(B)** Correlation plot comparing miRNA expression levels between PD synaptosome and PD cytosol. Each dot corresponds to an individual miRNA, with the diagonal line indicating equal expression between the two compartments. Deviations from the diagonal reflect differential distribution of miRNAs between the synaptosome and cytosol with red color intensity indicating level of significance. **(C)** Venn diagram illustrating the distribution of miRNAs between PD synaptosome and PD cytosol with significant overlap between the two compartments shaded in deep green color, and compartment specific expression of miRNAs, shaded in light green color for synaptosome and shaded in red color for cytosol. **(D)** Bar chart depicting the top 20 molecular function enrichment of miRNAs showing differential enrichment of miRNAs across molecular functions.

### MicroRNAs expression in PD soma versus UC soma

Next, we compared miRNA expression in cytosolic (soma) fractions of PD affected samples and non-PD affected samples. A total of 32 miRNA identified to be significantly deregulated in this cohort (Supplementary Table 3). These miRNAs can be found in hierarchical clustering and heatmap in Figure 5. Of the 32, none (0%) were altered at a p-value < 0.001, 1 at a p-value < 0.01, 31 at a p-value < 0.05. 8 (25%) were highly expressed. 18 (56.3%) were upregulated and 14 (43.8%) were downregulated as shown in Figure 5. Notably, miRNAs hsa-miR-224-5p_L1R-2, hsa-miR-486-3p, and hsa-miR-206 were significantly upregulated while hsa-mir-5585-p3_1ss7AD, and mmu-miR-6412_L-2R-1ss15AT were downregulated as shown in the volcano plot in Figure 6A. The correlation plot in Figure 6B showed similar expression of miRNAs in the PD cytosol and UC cytosol. Figure 6C showed a significant overlap of expression of miRNAs in PD cytosol and UC cytosol, but there was much less exclusive expression in PD cytosol compared to UC cytosol. In terms of molecular function enrichment, these miRNAs were significantly involved in protein binding and less involved in DNA-binding transcription activator activity, and RNA polymerase II specific binding (Figure 6D). Based on these observations, there appears to be less difference in expression of miRNA in PD cytosol vs UC cytosol than observed in the prior experimental cohorts. Out of the total miRNA pool, 1.86% showed a significant difference in the cytosol of PD affected individuals vs that of non-PD affected individuals. 1.05% were significantly upregulated and 0.82% were downregulated.

**Figure 5.**
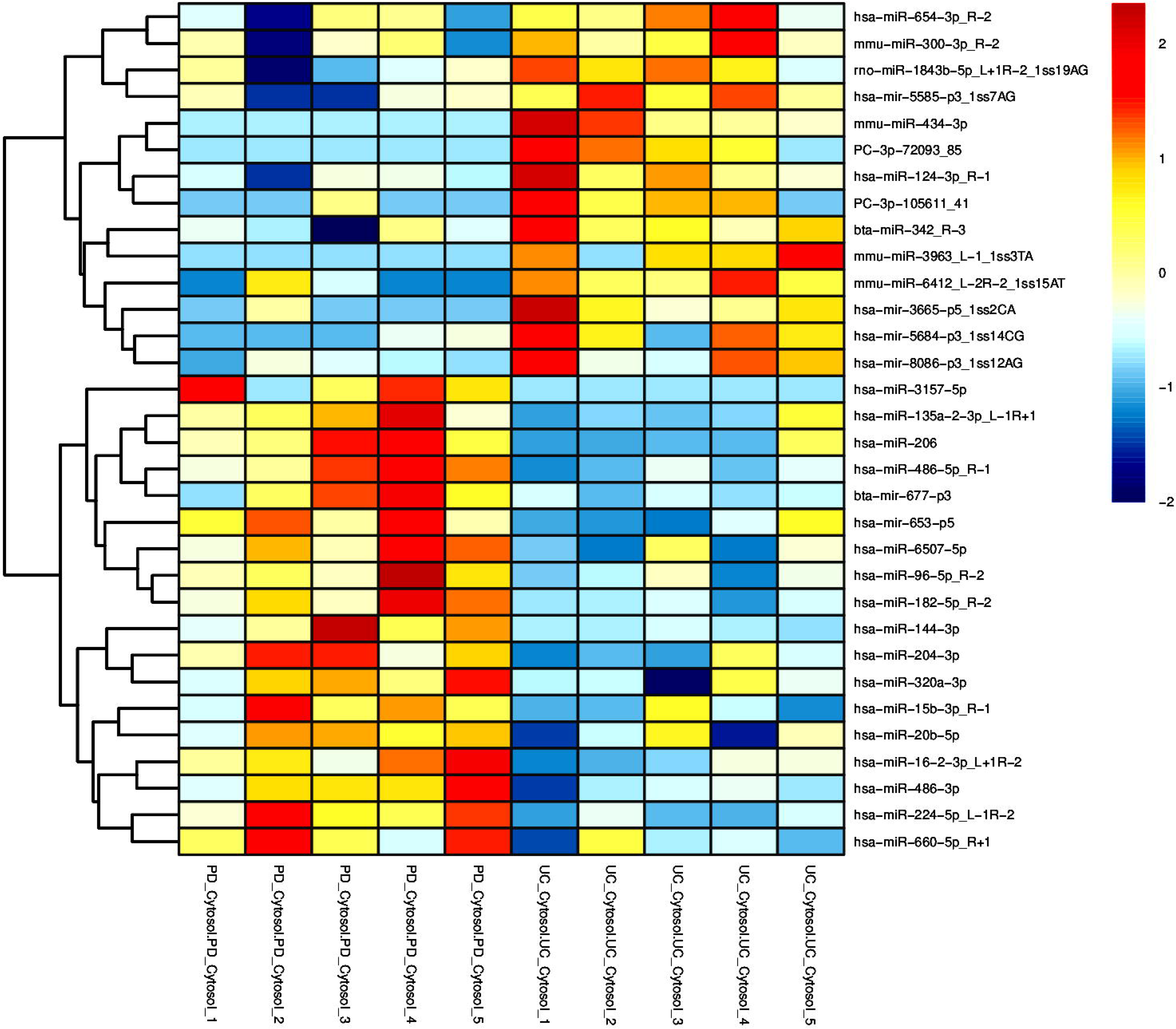
MiRNAs expression in PD cytosol vs UC cytosol. Hierarchical clustering and heatmap of 32 significantly distributed miRNAs in cytosol in PD vs UC samples. Red color intensity showed the miRNAs upregulation and blue color intensity showed the miRNAs downregulation.

**Figure 6.**
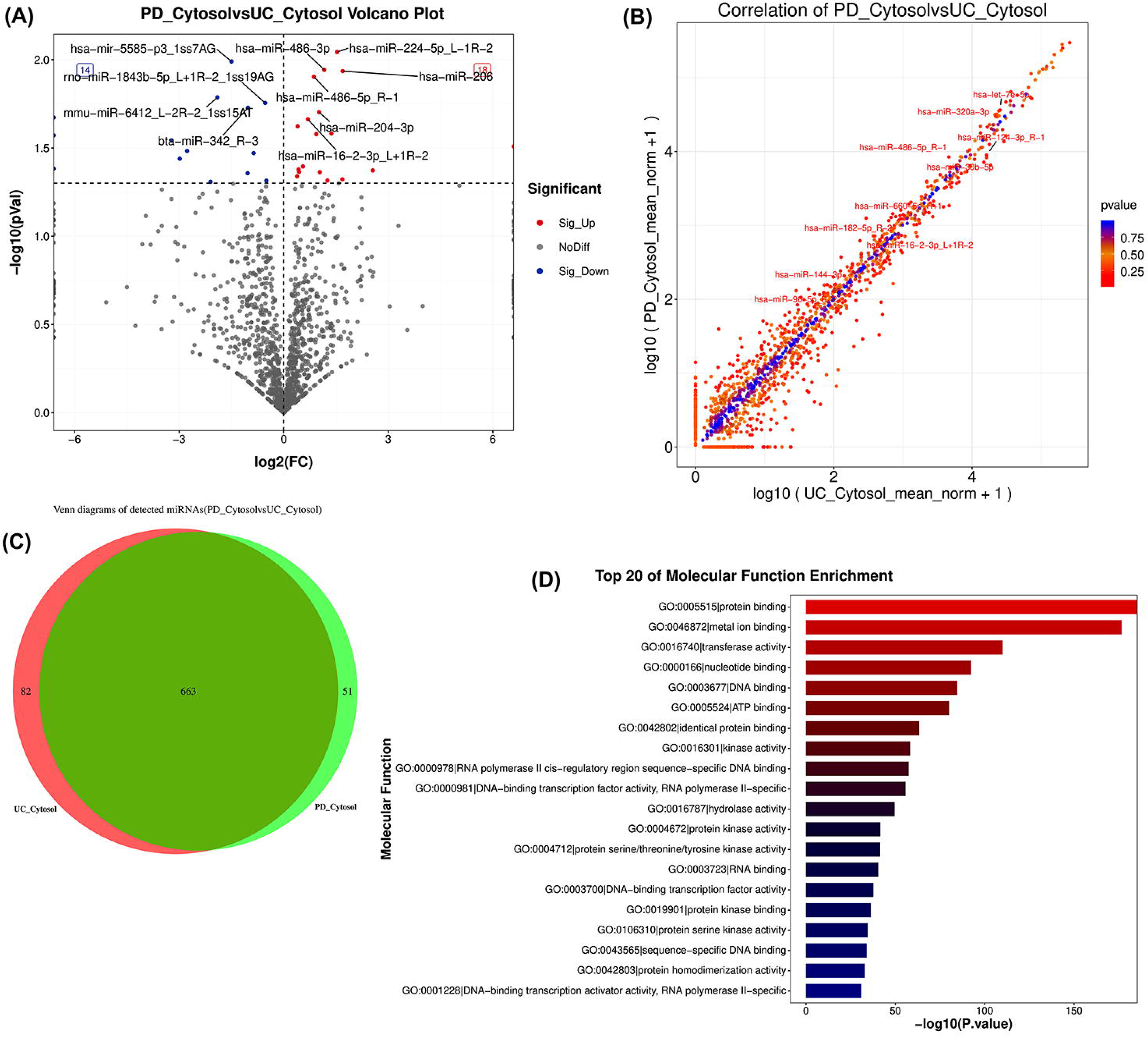
MiRNAs expression in PD cytosol versus UC cytosol. **(A)** Volcano plot depicting differentially expressed miRNAs in PD cytosol vs UC cytosol. Red dot showed the miRNAs upregulation, blue dot showed the miRNAs downregulation, and gray dot showed no difference. Significantly upregulated and downregulated miRNAs are highlighted. **(B)** Correlation plot comparing miRNA expression levels between PD cytosol and UC cytosol. Each dot corresponds to an individual miRNA, with the diagonal line indicating equal expression between the two cohorts. Deviations from the diagonal reflect differential distribution of miRNAs between the PD and UC samples with red color intensity indicating level of significance. **(C)** Venn diagram illustrating the distribution of miRNAs between PD cytosol and PD cytosol with significant overlap between the two cohorts shaded in deep green color, and cohort specific expression of miRNAs, shaded in light green color for PD cytosol and shaded in red color for UC cytosol. **(D)** Bar chart depicting the top 20 molecular function enrichment of miRNAs showing differential enrichment of miRNAs across molecular functions.

### MicroRNAs expression in PD synaptosome versus UC synaptosome

Lastly, we compared the microarray data for changes in miRNA expression between the synaptosomal compartment in PD-affected individuals vs non-PD-affected individuals. A total of 49 miRNA were found to be significantly deregulated in this cohort (Supplementary Table 4). These miRNAs can be seen in hierarchical clustering and heatmap in Figure 7. Of the 49, none (0%) were significantly altered at a p-value < 0.001, 6 at a p-value < 0.01, and 43 at a p-value < 0.05. Only 7 (14.3%) were highly expressed. 16 (32.7%) were upregulated and 33 (67.3%) were downregulated as shown in Figure 8a. Of note, miRNAs hsa-miR-6511b-5p, and hsa-miR-378g_R+2 was significantly upregulated while hsa-miR-1468-5p_R+1 and hsa-miR-9983-3b were downregulated as shown in the volcano plot in Figure 8A. The correlation plot in Figure 8B showed similar expression of miRNAs in the PD synaptosome and UC synaptosome. Figure 8C shows a significant overlap of expression of miRNAs in PD synaptosome and UC synaptosome, but there was much less exclusive expression in PD synaptosome compared to UC synaptosome. The distribution of miRNA in all cellular components was found to be highest in the cytosol and was relatively lower in protein containing complexes and neuron projections (Figure 8D). The most interesting finding is that all 7 of the miRNA found to be highly expressed were also determined to be downregulated in the PD-affected synaptosome fractions in comparison to the non-PD-affected synaptosome fractions. This suggests that the PD pathology may have some significant impact at the synapse. This finding warrants further investigation. Out of the total miRNA pool, only 1.4% are significantly deregulated in the synaptosomal fraction of samples from PD-affected individuals vs non-PD-affected individuals. Only 0.45% were upregulated and 0.92% were down regulated.

**Figure 7.**
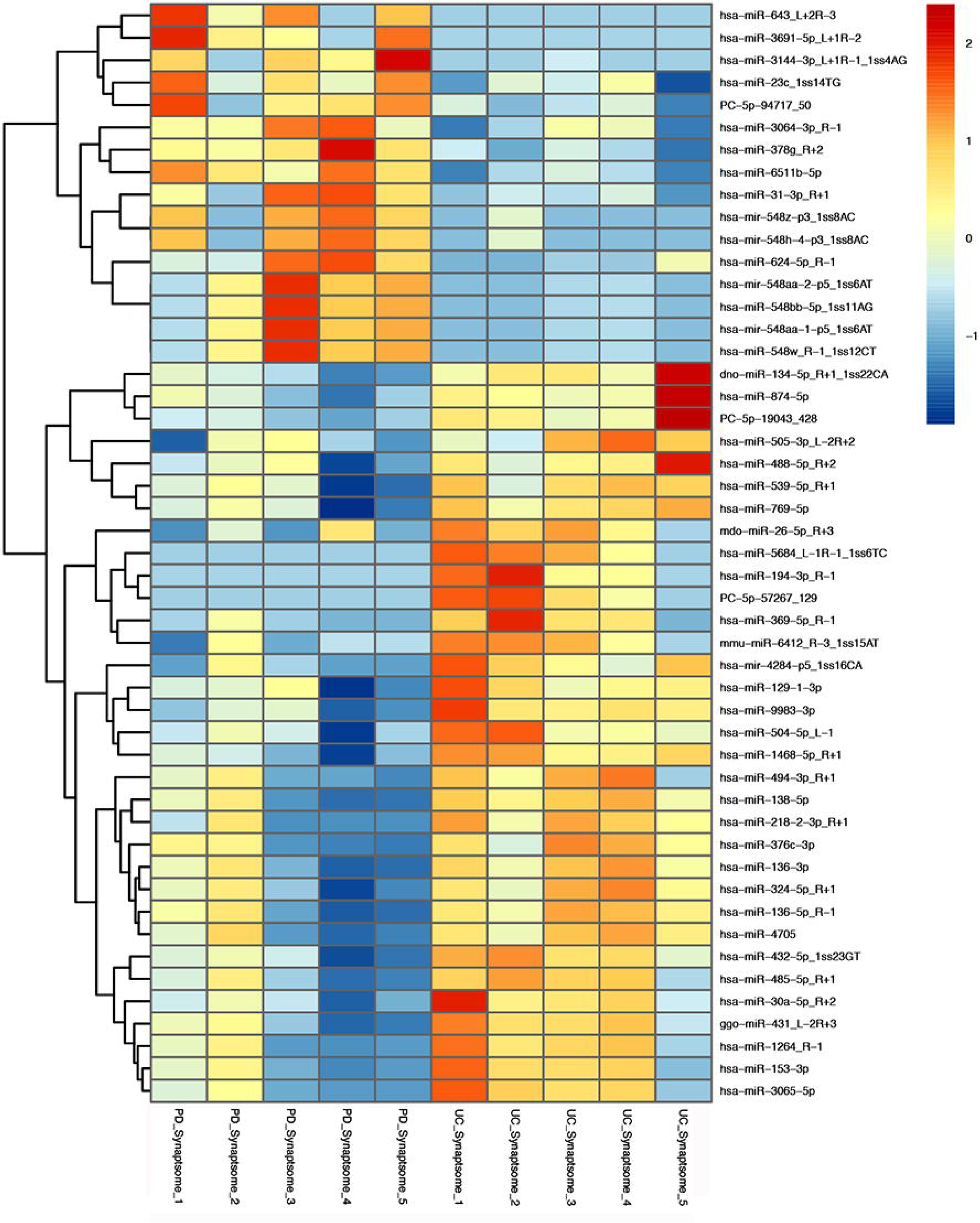
MiRNAs expression in PD synaptosome vs UC Synaptosome. Hierarchical clustering and heatmap of 49 significantly distributed miRNAs in synaptosomes in PD vs UC samples. Red color intensity showed the miRNAs upregulation and blue color intensity showed the miRNAs downregulation.

**Figure 8.**
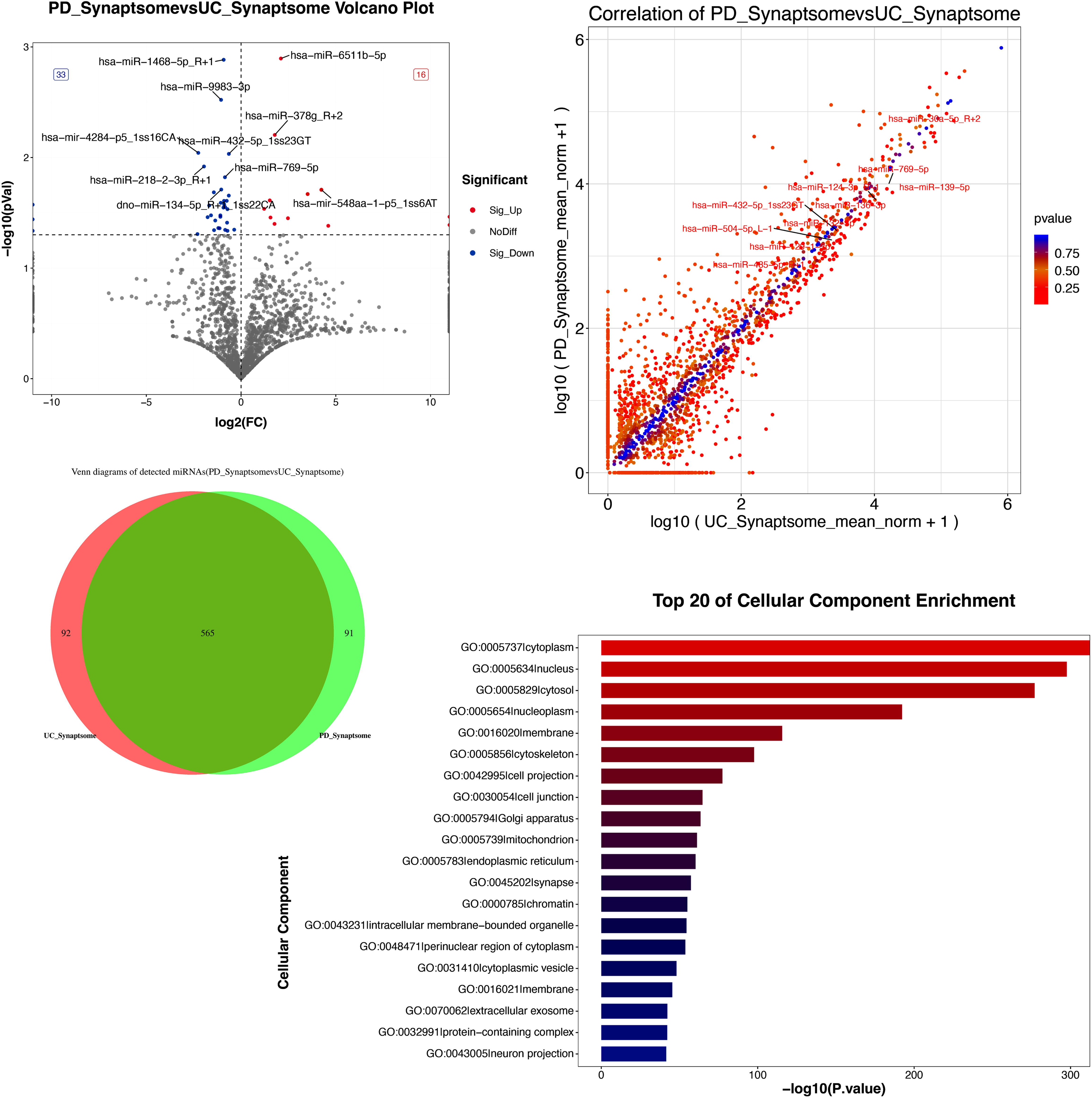
MiRNAs expression in PD synaptosome vs UC synaptosome. **(A)** Volcano plot depicting differentially expressed microRNAs in PD synaptosome vs UC synaptosome. Red dot showed the miRNAs upregulation, blue dot showed the miRNAs downregulation, and gray dot showed no difference. Significantly upregulated and downregulated microRNAs are highlighted. **(B)** Correlation plot comparing miRNA expression levels between PD synaptosome and UC synaptosome. Each dot corresponds to an individual microRNA, with the diagonal line indicating equal expression between the two cohorts. Deviations from the diagonal reflect differential distribution of miRNAs between the PD and UC samples with red color intensity indicating level of significance. **(C)** Venn diagram illustrating the distribution of miRNAs between PD synaptosome and PD synaptosome with significant overlap between the two cohorts shaded in deep green color, and cohort specific expression of miRNAs, shaded in light green color for PD synaptosome and shaded in red color for UC synaptosome. **(D)** Bar chart depicting the top 20 cellular component localization of miRNAs showing differential enrichment of miRNAs across cellular regions.

## DISCUSSION

Multiple studies have demonstrated that miRNAs influence various synaptic functions (Edbauer et al., 2010; Lee et al., 2016; Tsujimura et al., 2016; Wu et al., 2018; McNeill et al., 2020). While it is well established that synaptic miRNAs contribute to various synaptic functions, the specific role of synaptosome-associated miRNAs in the pathogenesis of PD remains unclear. Therefore, we hypothesized that synaptosomal miRNA expression is altered in PD. For the first time, we provide a comprehensive analysis of miRNA expression in synaptosomal and cytosolic fractions of the substantia nigra from post-mortem brains of PD patients and UC.

So, how does this information benefit the academic and medical community? It is known that there is a strong association between destruction of dopaminergic neurons in the substantia nigra and the development of PD. Anatomically, most of these neurons originate in the SNpc. Substantia nigra pars reticulata (SNpr) neurons are primarily gabaergic, and do undergo some pathological changes in PD, such as increased activity due to the depletion of dopamine in the striatum. However, pathological changes are not to the degree seen in the SNpc. We have essentially shown that there is a significant change to the miRNA profile in PD which is a representation of the damages or changes that occur in the substantia nigra. In the future, we can explore the effects of altering the expression of key miRNA in this region, to observe if it has any positive, negative, preventative, deleterious, or curative effects in PD pathology.

Furthermore, we analyzed the synaptosomal miRNA in this region. Referring back to the anatomy, we have essentially acquired the miRNA profile in the synapses of gabaergic neurons originating in the globus pallidus and striatum which terminate in the SNpc and glutamatergic neurons originating in the subthalamic nucleus which terminate in the SNpc. This subset of the information we acquired may prove to be valuable and we strongly encourage further research into the concepts. Nonetheless, what may be more useful is to employ these methods using tissue from the striatum (caudate-putamen). Synaptosome extraction and miRNA analysis of the striatum, would give us the miRNA profile of synapses of dopaminergic neurons originating in the SNPc and terminating in the caudate and putamen. Doing so, would give the academic and medical community a wealth of information regarding synaptic function and synaptic dysfunction in PD, regulation of alpha-synucleinopathy, as well as open an avenue for pharmacotherapeutic strategies that actually target the progression of PD. Our lab is actively pursuing elucidating this information.

It should be noted, as depicted in the pie chart analyses in figures 2,4,6, and 8, that > 70% of miRNA that are expressed in the synapse are also expressed in the cytosolic fractions. Only a small subset of miRNAs is distinctly found in the synapse. Additionally, the majority of the miRNA population did not show any significant changes in the synaptosome and cytosol. Only a small fraction of miRNA showed significant changes among synaptosomes and cytosol. These findings support a conclusion that miRNAs are uniformly distributed within the neuron with a relatively small fraction being localized to the synapse. These synaptosomal miRNA are either synthesized locally at the synapse or are transported from different compartments, such as the soma, to the synapse. Additional research into this process is warranted.

When comparing the miRNA within the cytosolic fraction in PD affected samples to the unaffected control samples, we found a group of miRNAs that are worth highlighting. Hsa-miR-224-5p, hsa-miR-204-3p and hsa-miR-144-3p which usually merits average expression is upregulated in PD, while hsa-miR-124-3p was downregulated in PD. Researchers have shown that this miRNA can be manipulated, by way of inhibition, to significantly improve neuron survival and reduce neuronal apoptosis in a model of oxygen-glucose deprivation (Liu et al. 2020). Interestingly, we found hsa-miR-124-3p, a miRNA usually found to be highly expressed, is downregulated in PD. miR-124-3p has been shown to have neuroprotective effects in PD, as it reduced neurotoxicity in the 6-hydroxydopamine (6-OHDA)-induced model of PD (Esteves et al., 2022). Further research into the effect of manipulating these miRNAs in PD is warranted and strongly encouraged.

In summary, our study identified the soma (cytosolic) and synaptosomal miRNAs that are deregulated in PD. However, our findings in this study are limited by non-use of striatal tissue, and therefore there is a void of the miRNA profile of the synapses of the dopaminergic neurons involved in PD. Our ongoing research addresses this void and seeks to further investigate the underlying molecular mechanisms of the miRNA that play an important role in PD.

## ACKNOWLEDGEMENTS

This research was funded by the National Institute on Aging (NIA), National Institutes of Health (NIH), grant number K99AG065645, R00AG065645, R00AG065645-04S1, SARP mini grants TTUHSC EP, Edward N. & Margaret G. Marsh Foundation and TTUHSC EP MTM Startup Funds to S.K.

## AUTHOR CONTRIBUTIONS

Conceptualization and supervision: SK; experimental performance: SK, MO, GG, BS, PZ, and DR; analysis, interpretation, and validation of data: SK, MO, and BS; writing and original draft preparation: MO, SK and PZ; review, editing, and finalization of manuscript: SK, MO and RL. All authors have read and agreed to the published version of the manuscript.

## DATA AVAILABILITY

Data will be available upon request.

## COMPETING INTERESTS

None

## Notes

### Competing Interest Statement

The authors have declared no competing interest.

