## Supplementary figures and images for "MicroRNAs alteration and unique distribution in the soma and synapses of substantia nigra in Parkinson’s disease"

### Supplementary Figure 1

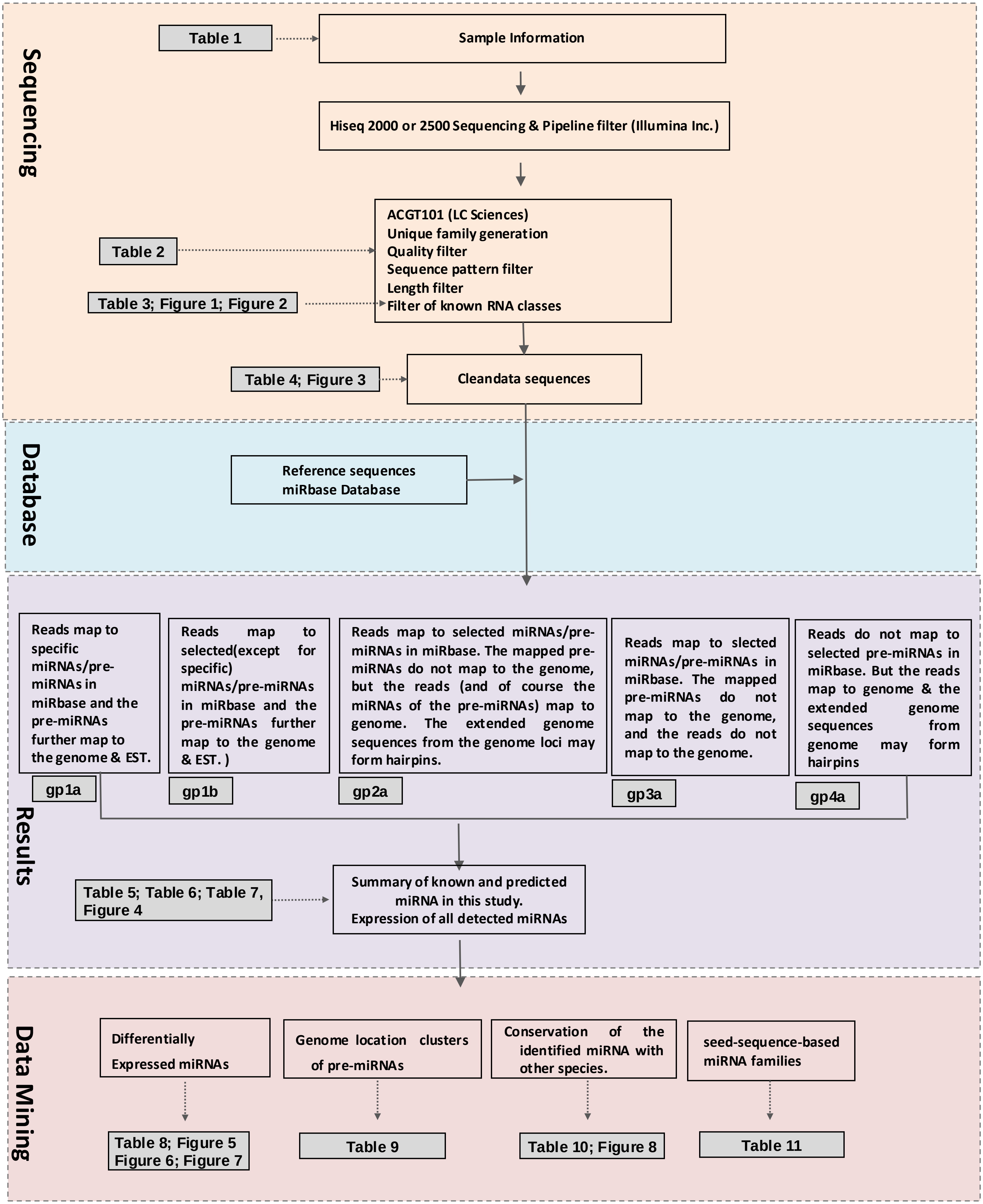


**Figure S1. Analysis workflow and the corresponding tables & figures.**
